# Immunogenicity of different types of adjuvants in bacterial vaccine

**DOI:** 10.64898/2025.12.28.696776

**Authors:** Yue-Yue Li, Nan Wang, Qiang Han, Teng Feng, Zhuang Zhu, Teng-Lin Xu, Ke-Ji Quan, George Fei Zhang

**Affiliations:** International Joint Research Center for Microbiology and Immunology, Wohua Bioeng, Binzhou, Shandong, 256600; Institute of Basic Medicine, North Sichuan Medical College, Nanchong, Sichuan, China; Department of Molecular Biology, Umeå University, Umeå, Sweden

**Keywords:** Different Adjuvant, Commercial vaccine, Avibacterium paragallinarum, Humoral immune, mineral oil adjuvant, Aluminum adjuvant

## Abstract

This study aims to investigate four common trivalent avian infectious coryza (AIC) vaccines available on the market. It detects the antibody levels of three serotypes at 7 days and 14 days after the first immunization, and at 30 days, 90 days, and 180 days after the second immunization using HI (Hemagglutination Inhibition Test) and ELISA (Enzyme-Linked Immunosorbent Assay). Additionally, the levels of cytokines including IFN-γ (Interferon-Gamma), IL-2 (Interleukin-2), and IL-3 (Interleukin-3) are measured. Furthermore, the protective rates of four of these vaccines are verified 180 days later. It demonstrated that the immunogenicity of mineral oil adjuvanted vaccine is higher than that of Aluminum adjuvanted. The animals should be boosted with another booster immunization six month post the first immunization to protect them from infectious coryza.

## 1 Introduction

Avian infectious coryza is an acute upper respiratory infectious disease caused by *Haemophilus paragallinarum*. This disease leads to stunted growth in chickens and a decrease in egg production rate, inflicting significant economic losses on the poultry industry(1). Given the high incidence and harmfulness of this disease, vaccination has become an important measure for preventing avian infectious coryza.

Currently, there is a wide variety of avian infectious coryza vaccines available on the market, mainly including different types such as monovalent vaccines, bivalent vaccines, trivalent vaccines, bivalent combined vaccines, and trivalent combined vaccines. These vaccines differ in terms of serotype coverage, immunogenicity, protective efficacy, safety, and applicable chicken flocks(2). For instance, monovalent vaccines target a single serotype of Haemophilus paragallinarum, while bivalent and trivalent vaccines can prevent infections caused by two or three different serotypes simultaneously, enabling more comprehensive response to complex epidemic strains(3).

Vaccines of the same type produced by different manufacturers vary in production processes, adjuvant selection, and quality control measures, which in turn affect the vaccines’ immunological efficacy and practical application effects. For example, oil-emulsion inactivated vaccines and aluminum hydroxide gel-adjuvanted inactivated vaccines each have their own advantages and disadvantages in terms of the type of immune response induced, the duration of antibody level maintenance, and the impact on the egg production rate of laying hen flocks.

With the large-scale development of the poultry industry, farmers and veterinarians have become increasingly cautious in the selection of avian infectious coryza vaccines. This study selected two common types of vaccines on the market—oil-emulsion vaccines and aluminum hydroxide gel-adjuvanted vaccines—for comparison, aiming to explore the immunogenicity and antibody persistence duration of the two types of vaccines.

## 2 Materials and Methods

### 2.1 Bacteria and study design

The 42-day-old specific pathogen-free (SPF) chickens were sourced from Binzhou Wohua Biotechnology. A total of 68 chickens were randomly divided into 5 groups, with 10 chickens in each group, and 8 chickens in the control group. APG vaccines were purchased from both domestic and international sources on the market, with specific information shown in Table 1. The immunization route was subcutaneous injection in the neck, and the immunization dose was 0.5 mL per chicken.

**Table 1.** Information of Vaccines.

| Vaccine | Company | Description |
| --- | --- | --- |
| Vac-1 | New Jiuzhou (Kitasato) | Bivalent Inactivated Vaccine for Avian Infectious Coryza |
| Vac-2 | Beijing Aifeikes Biological Technology Co., Ltd. | Trivalent Vaccine for Avian Infectious Coryza |
| Vac-3 | Merck Animal Health | Trivalent Inactivated Vaccine for Avian Infectious Coryza |
| Vac-4 | Beijing Xinde Weite Technology Co., Ltd. | Trivalent Inactivated Vaccine for Avian Infectious Coryza (Serotype A + B + C) |

### 2.2 Reagents

The Chicken Infectious Coryza IgG Antibody ELISA Detection Kit was self - made by our company; Chicken IL - 2 ELISA Detection Kit (Nanjing Senbeijia Biotechnology), Chicken IL - 6 ELISA Detection Kit (Nanjing Senbeijia Biotechnology), and Chicken IFN - γ ELISA Detection Kit (Xinbosheng Biotechnology) were used. Other reagents included glutaraldehyde (Kemiou Chemical Reagent), thimerosal (Sinopharm Group), potassium dihydrogen phosphate (Sinopharm Group), and dipotassium hydrogen phosphate (Tianjin Tianli Chemical Reagent).

### 2.3 Efficacy Determination

42-day-old SPF chickens were used for vaccine efficacy testing. The specific grouping and immunization details are shown in Table 2. The second immunization was conducted 14 days after the first immunization. The challenge strain was APG - 91, and the incidence was observed for 7 days after the challenge.

**Table 2.** Grouping and Immunization for Efficacy Testing.

| Vaccine | Company | Immunization Dose | Number of Chickens | Challenge Strain | Challenge Dose |
| --- | --- | --- | --- | --- | --- |
| Vac-1 | Kitasato | 0.5 mL/bird | 10 | APG - 91 | 4000 CFU |
| Vac-2 | Meidian | 0.5 mL/bird | 10 | APG - 91 | 4000 CFU |
| Vac-3 | Merck (Avian Biwei) | 0.5 mL/bird | 10 | APG - 91 | 4000 CFU |
| Vac-4 | Xinde (Biduo Bao) | 0.5 mL/bird | 10 | APG - 91 | 4000 CFU |
| - | Control | 0.5 mL/bird | 8 | APG - 91 | 4000 CFU |
| - | Total | - | 68 | - | - |

### 2.4 Detection of Specific Antibodies

Blood samples were collected to separate serum before immunization, at 7 days and 14 days after the first immunization, and at 30 days, 90 days, and 180 days after the second immunization for the determination of specific IgG antibody levels against infectious coryza in chickens using ELISA.

### 2.5 Determination of HI Hemagglutination Titer

Blood samples were collected to separate serum at 7 days and 14 days after the first immunization, and at 30 days, 90 days, and 180 days after the second immunization for the determination of hemagglutination titer.

### 2.6 Detection of Inflammatory Cytokines

Blood samples were collected to separate serum at 7 days and 14 days after the first immunization. According to the manufacturer’s instructions, ELISA kits were used to detect the protein contents of inflammatory cytokines in the serum, including chicken IL-6, IL-2, and IFN-γ.

## 3 Results

### 3.1 Results of Vaccine Efficacy Determination

The challenge with APG-91 was conducted 180 days after the two immunizations, and the chickens after immunization were observed continuously for 7 days after the challenge. The challenge protection results are shown in Figure 1. Xinliubi vaccine showed a good protection rate, reaching 100% protection. Kitasato and Meidian, as aluminum hydroxide - adjuvanted vaccines, had a short immune duration with a protection rate of less than 20%. Xinde and Merck oil - adjuvanted vaccines against infectious coryza in chickens achieved a protection rate of more than 70% after the challenge.

**Figure 1.**
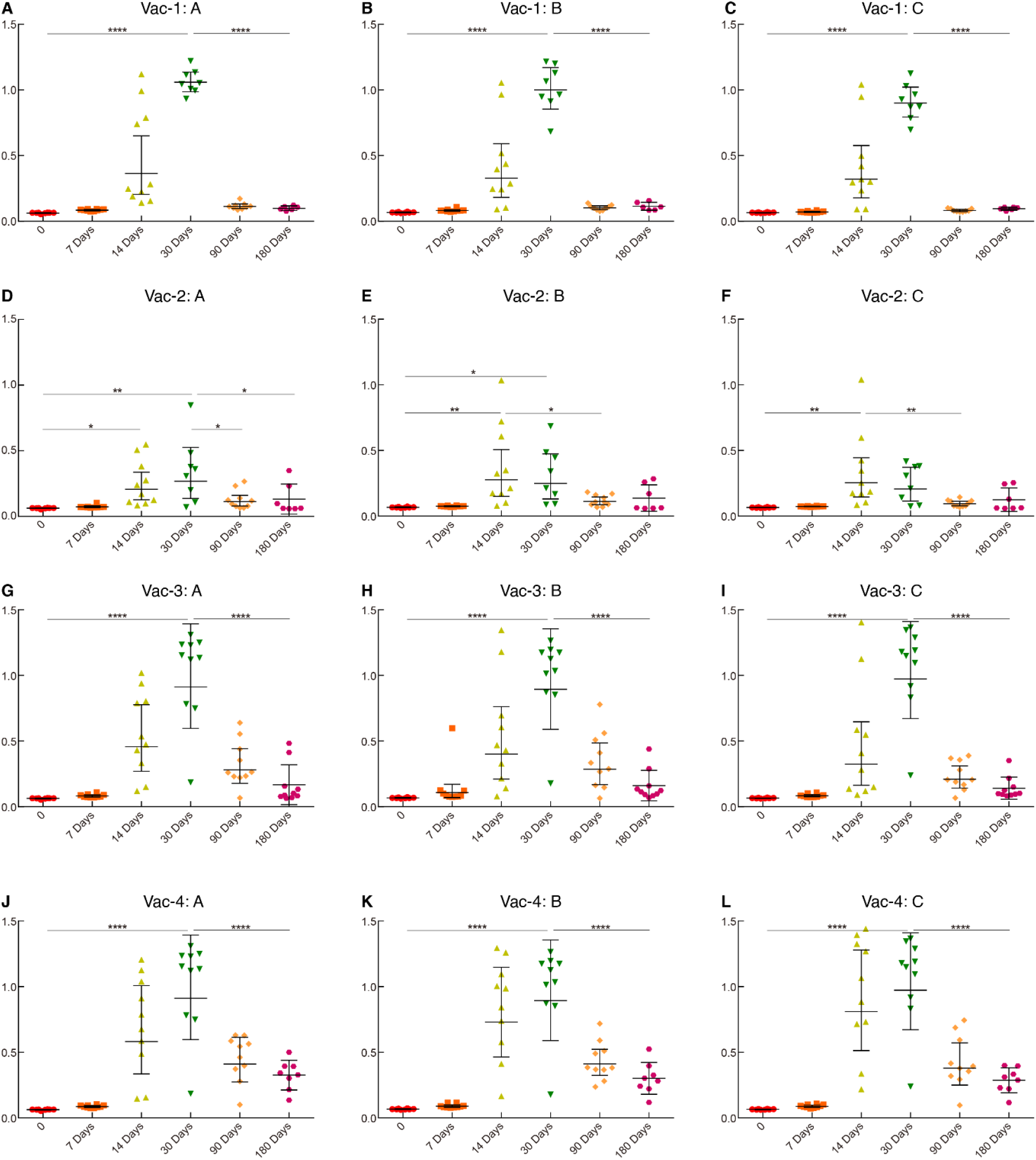

### 3.2 Detection of Specific Antibodies

Blood samples were collected to separate serum before immunization, at 7 days and 14 days after the first immunization, and at 30 days, 90 days, and 180 days after the second immunization. The level of specific IgG antibodies against infectious coryza in chickens was determined by ELISA, and the results are shown in Figure 2. The antibody levels of all vaccines began to increase after the first immunization, reached a peak one month after the second immunization, started to decrease at three months, and continued to decrease at six months. Among them, the Xinliubi vaccine induced antibody production earlier and had a longer duration of antibody persistence.

**Figure 2.**
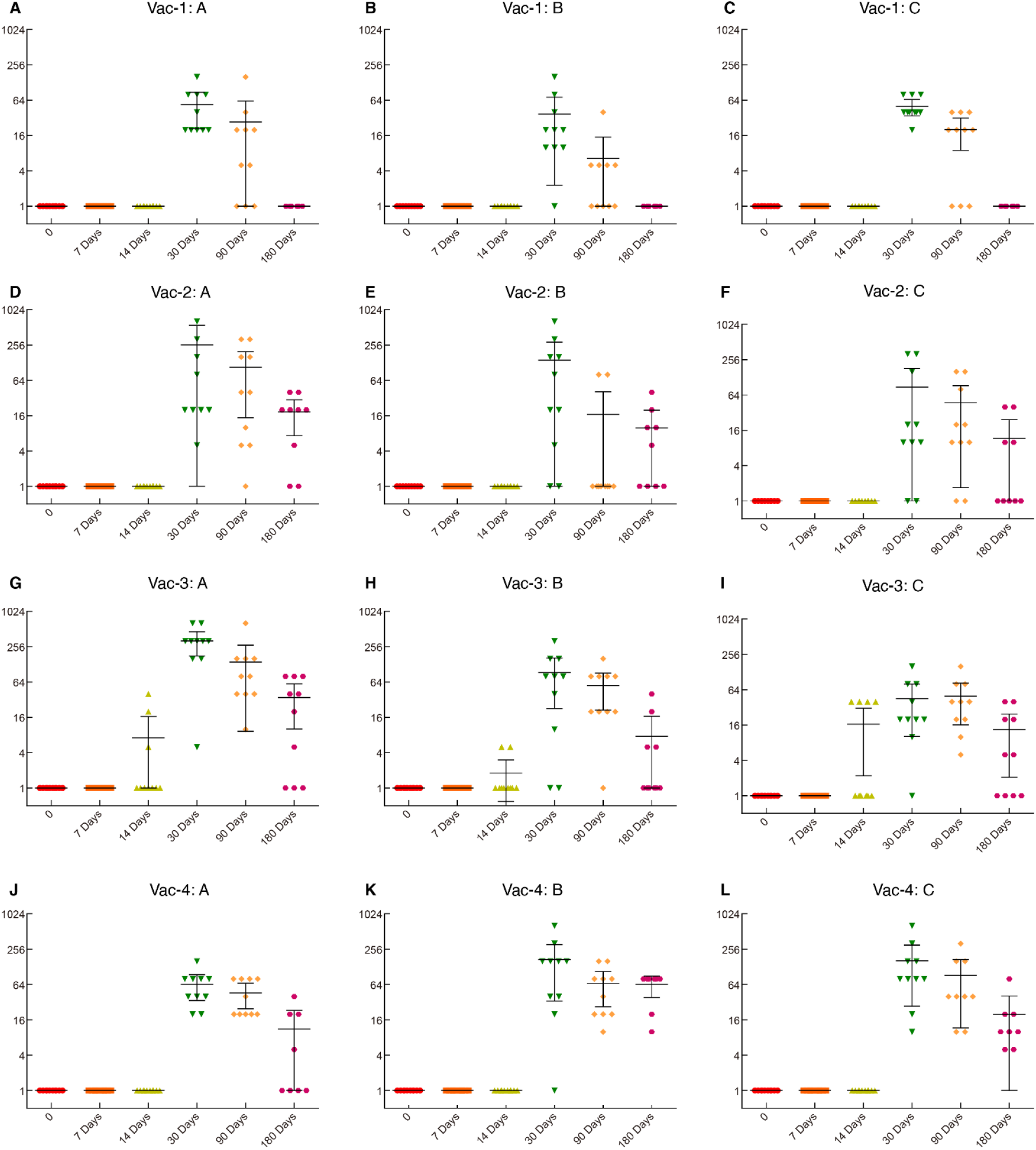
Vaccine Antibody Levels.

**Figure 3.**
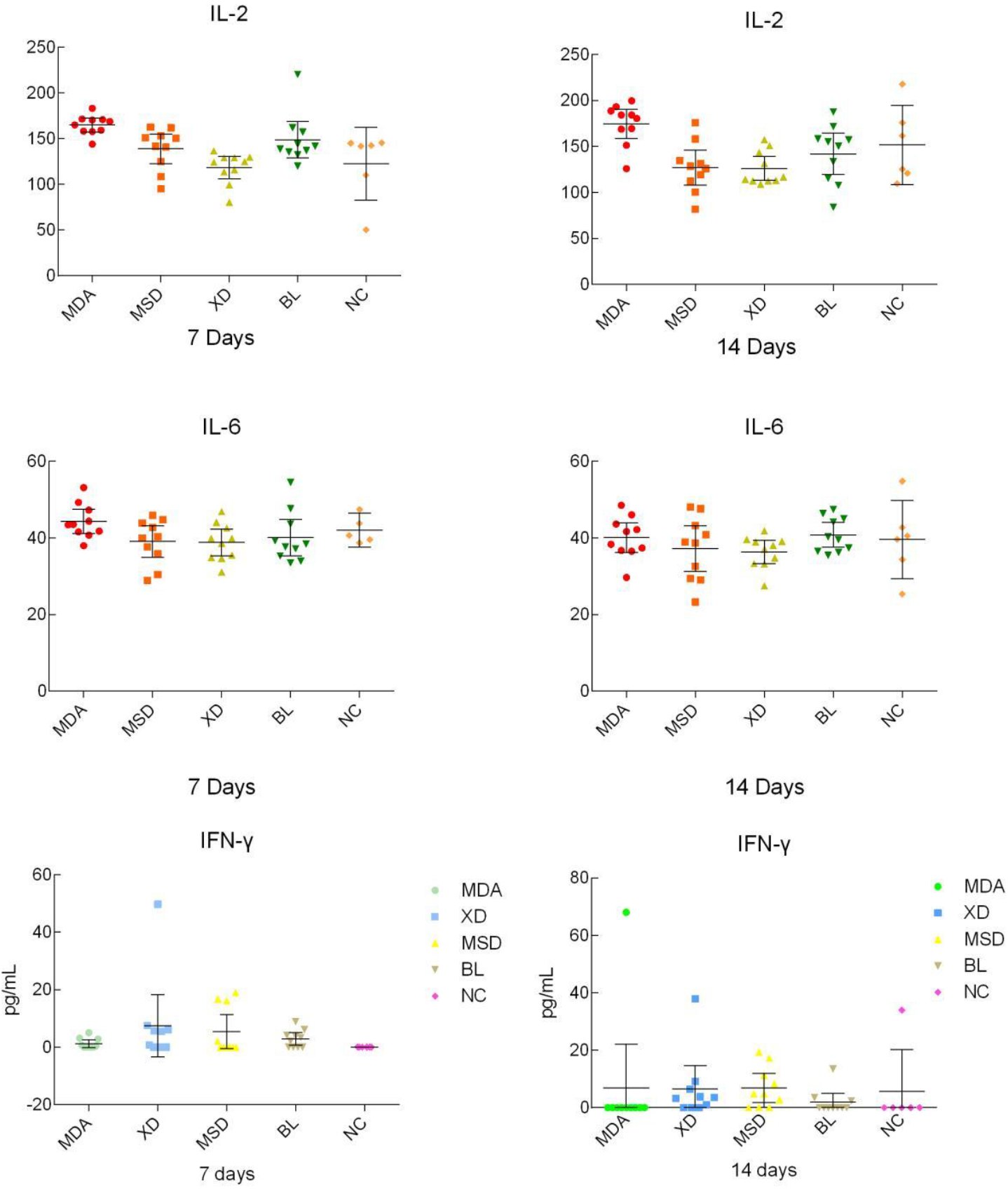

**Figure 4.**
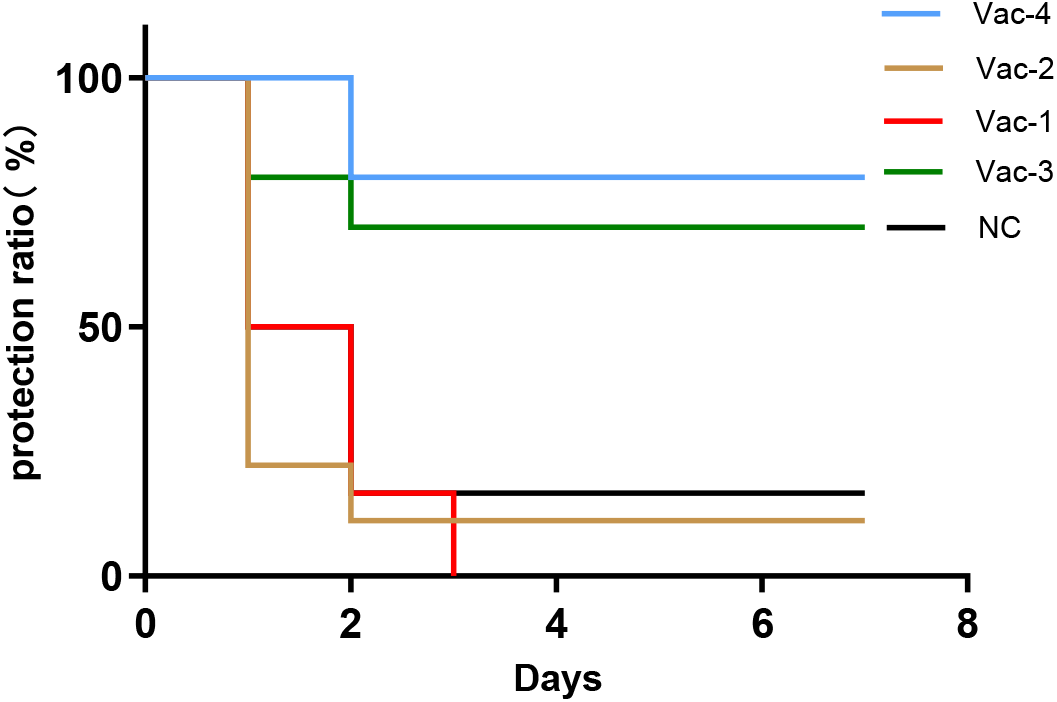
Vaccine Protection Status.

### 3.3 Detection of HI Antibodies

HI tests were performed to detect the three serotypes of infectious coryza in chickens at five time points: 7 days, 14 days, 30 days, 90 days, and 180 days after immunization. No HI titer was detected for Xinliubi vaccine. Antibodies of Merck oil-adjuvanted vaccine began to appear at 14 days. Except for Xinliubi vaccine, all other vaccines reached the antibody peak one month after immunization, which was consistent with the ELISA results. The antibody levels decreased slightly at 90 days and continued to decrease at 180 days.

### 3.4 Detection of Cytokines

Serum samples collected at 7 days and 14 days after immunization were used for cytokine detection. The levels of cytokines IL-2 and IL-6 in Meidian vaccine were significantly higher than those in other vaccines at 7 days and 14 days.

## 4 Discussion

In this experiment, efficacy determination, specific antibody detection, HI antibody detection, and cytokine detection were conducted on different vaccines against infectious coryza in chickens, and important results were obtained, which are discussed in depth as follows(4). In contrast, Hokuriku and Meidian aluminum hydroxide gel-adjuvanted vaccines had a short immune duration with a protective rate of less than 20%, which may be due to the rapid release of antigens, failing to maintain immune stimulation. Xinde and Merck oil-adjuvanted vaccines had a protective rate of more than 70%.

The detection of specific antibodies showed that the antibody levels of all vaccines increased after the first immunization, reached a peak one month after the second immunization, and then gradually decreased(5). Xinliubi vaccine had an earlier onset of antibody production and a longer duration of antibody persistence, which was consistent with its high protective rate, indicating that it can trigger the immune response earlier and maintain a high antibody level. The changes in antibody levels reflect the dynamics of immune memory. The antibody peak after the second immunization may be due to the activation of memory B cells by the second immunization, resulting in the production of a large number of antibodies.

The HI antibody detection results were similar to those of specific antibody detection. No HI titer was detected for Xinliubi vaccine, which may be due to its different immune mechanism that does not rely on HI antibodies(6). Antibodies of Merck oil-adjuvanted vaccine began to appear at 14 days, and the antibodies of other vaccines reached the peak one month after immunization. The antibody levels decreased slightly at 90 days and continued to decrease at 180 days, which was consistent with the ELISA results, confirming the regularity of the immune response(7).

In conclusion, different vaccines against infectious coryza in chickens show significant differences in immune efficacy, antibody levels, and cytokine regulation(8). The Xinliubi vaccine has prominent advantages and great application potential; the Xinde and Merck Sharp & Dohme (MSD) oil-adjuvanted vaccines have certain immune effects and can be optimized; the Hokuriku and Meidian aluminum hydroxide gel-adjuvanted vaccines need to have their formulas or immunization procedures improved; and in-depth research on the immune mechanism of Meidian vaccines should be conducted(9). Future studies can explore immune mechanisms, optimize formulas and procedures, and enhance the immune efficacy and protection rate of vaccines(10).

Significant differences in vaccine efficacy: The challenge protection rate of the Xinliubi vaccine reaches 100%, and it still maintains good efficacy 180 days after immunization, showing great application potential. The Hokuriku and Meidian aluminum hydroxide gel-adjuvanted vaccines have a short immune duration and low protection rate, so their formulas or immunization procedures need to be optimized(11). The Xinde and Merck Sharp & Dohme (MSD) oil-adjuvanted vaccines have a certain level of protective capacity, but there is still room for improvement compared with the Xinliubi vaccine.

Characteristics of antibody detection results: The antibody levels of all vaccines first increase and then decrease. The Xinliubi vaccine produces antibodies early and maintains them for a long time, which is related to its high protection rate. Hemagglutination Inhibition (HI) antibody detection shows that the immune mechanism of the Xinliubi vaccine may be different, as it does not mainly rely on HI antibodies(12).

Cytokines and future research: The Meidian vaccine can increase the level of immunomodulatory cytokines and activate immune cells in the early stage, but its protection rate is low, indicating that the protective effect of vaccines is affected by multiple factors. In the future, it is necessary to conduct in-depth research on the immune mechanisms of different vaccines, optimize their formulas and procedures, and develop more efficient and long-lasting vaccines to meet the needs of disease prevention and control.

## Acknowledgment

This study was funded by the Taishan Industrial Expert Programme of Shandong Province (NO. tscx202306107). This study was supported by the Sichuan Natural Science Foundation (2025ZNSFSC1084).

## Author’s contribution

GF Zhang conceived and designed the study. YY Li, N Wang, Q Han, and T Feng executed the experiment. GF Zhang processed data analysis and writing. GF Zhang, KJ Quan, TL Xu, and Z Zhu edited the manuscript. All authors approved the final version of the manuscript.

## Data availability statement

The raw data supporting the conclusions of this article are available from the authors upon reasonable request.

